# A fast immune priming that confers a complete infection resistance on silkworm (*Bombyx mori*)

**DOI:** 10.1101/205369

**Authors:** Atsushi Miyashita

**Affiliations:** Department of Psychology and Neuroscience, Dalhousie University, 1355 Oxford Street, Halifax, NS B3H 4R2, Canada

**Keywords:** immune priming, silkworm, collagen, *Escherichia coli*, *Pseudomonas aeruginosa*

## Abstract

I have previously reported an immune priming in silkworm triggered by peptidoglycans, which is long-lasting but slow-acting. Here I report a faster immune priming, which can be triggered by an injection of gelatin or collagen, that confers a complete infection resistance to Gram-negative bacteria on silkworms. Gelatin-injected silkworms showed 100% viability in a lethal dose of a Gram-negative bacterial infection within two hours after the gelatin injection. Injection of collagen showed a similar effect. Whereas, an injection of non-gelatin protein (bovine serum albumin) solution did not induce such reaction. These results suggest that the silkworm possesses a fast and gelatin-inducible pathway that confers infection resistance to Gram-negative bacteria, which may act as a front-line defense. This finding highlights the potency of gelatin as a tool for investigating the primed immune responses in insect species.

## 1. Introduction

I have previously reported that a pre-injection of heat-killed cells of Gram-negative bacteria, or its cell wall moiety (peptidoglycans), confers infection resistance to otherwise lethal bacterial infection in silkworm (Miyashita et al., 2014; Miyashita et al., 2015). Although the peptidoglycan-inducible immune priming conferred a full resistance to pathogenic Gram-negative bacteria, its proposed molecular mechanism (i.e. antimicrobial peptide upregulation) takes long time to activate (Miyashita et al., 2014), which thus is not sufficient to suppress bacterial infection ongoing.

Apart from antimicrobial peptides, invertebrate species possess faster immune responses such as hemocytes activation (Johansson et al., 2000) or melanization (Jiravanichpaisal et al., 2006; Johansson and Soderhall, 1989). These pathways may act as a front-line defense system protecting host animal from wide range of pathogens, which, however, are not pathogen selective but rather attack any xenobiotic components regardless of its biological/chemical properties. In insects, immune system has been shown to self-damage (Sadd and Siva-Jothy, 2006). It is, therefore, plausible that an over activation of such immune response may lead to negative impacts on the animal’s life history traits (Pursall and Rolff, 2011). Given these fact, it is somewhat reasonable to assume a type of front-line defense system that is more pathogen-selective.

To find such a front-line defense system, I searched for substances that can confer infection resistance in the silkworm faster than peptidoglycans do. As a result, a pre-injection of indian-ink (a mixture solution of carbon particles and animal glues) was found to confer infection resistance to Gram-negative enterohemorrhagic *Escherichia coli* O-157 (data not shown) on silkworms. Whereas, the indian-ink did not show immune priming effect against Gram-positive bacteria (*Staphylococcus aureus*). The indian-ink was then fractionated into supernatant and precipitate by ultracentrifugation, and the supernatant fraction showed the priming effect (data not shown). Because the major component of the soluble fraction (i.e. supernatant) of indian-ink is gelatin, I examined whether gelatin exhibits immune-priming effect in silkworm. I also demonstrate, in this article, that it confers a full infection resistance to Gram-negative bacteria within two hours, and that the system is not triggered by non-gelatin protein.

## 2. Materials and methods

### 2.1 Silkworm

Silkworms were reared as described previously (Miyashita et al., 2012). Larvae on day 2 in 5^th^ instar were used in the experiments.

### 2.2 Immune priming and infection experiment

Immune priming experiments and infection experiments were performed as described in the previous report (Miyashita et al., 2014). For infection experiment, cultured live cells of a pathogenic *Escherichia coli* (O-157:H7 Sakai strain) which was previously shown to be lethal to the silkworm (Miyashita et al., 2012), was suspended in saline, and injected in hemocoel of the silkworm, as previously reported (Miyashita et al., 2012; Miyashita et al., 2014).

### 2.3 Reagent

Gelatin (Sigma, Lot #117F0197) and bovine serum albumin (BSA) (Sigma, Lot #104K0721) was dissolved in sterile saline (containing 0.90% sodium chloride). The saline was prepared as previously describe (Miyashita et al., 2014).

## 3. Results and Discussion

The silkworm pre-injected with gelatin solution showed infection resistance to O-157 (enterohemorrhagic *E. coll*), whereas the same amount of bovine serum albumin (BSA) did not show such effect (Fig. 1). The ED_50_ value of gelatin was 1.7 μg/larva, while that of BSA was greater than 250 μg/larva. Injection of a gelatin component, collagen, also conferred infection resistance to a Gram-negative bacterium (*Pseudomonas aeruginosa*) on silkworm (Fig. S1), and both porcine and bovine collagen exhibited the immune priming effect (Fig. S1). These results indicate that the silkworm reacts to gelatin and collagens, which triggers an immune priming to Gram-negative bacteria. Also, the result suggests that the reaction is not triggered universally by any protein molecules but rather specific to gelatin components.

**Fig. 1.**
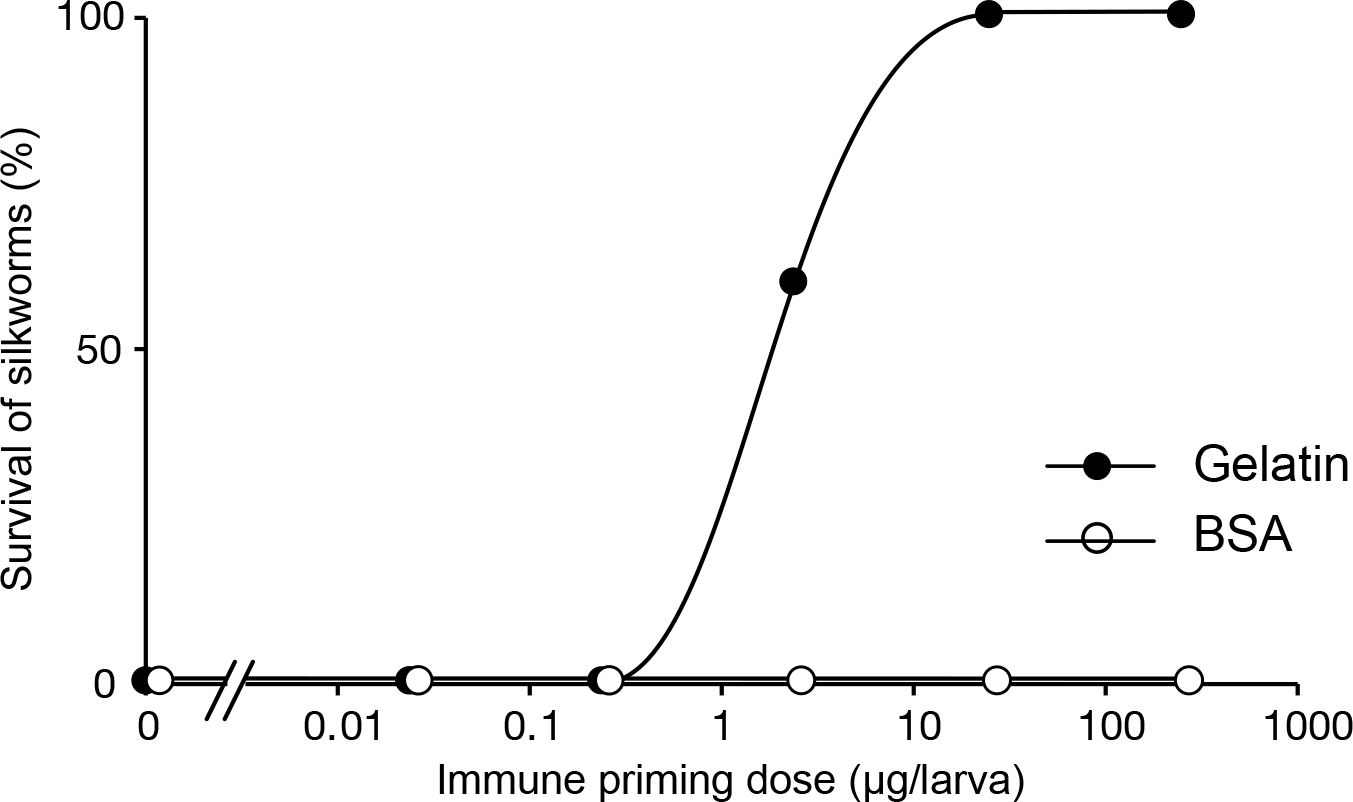
Immune priming effect of gelatin on silkworm. Silkworms were first injected with different doses of gelatin or bovine serum albumin (BSA), incubated for 4 hours at 27°C,and injected live cells of pathogenic *E. coli* (O-157:H7) as previously reported (Miyashita et al., 2012). The vertical axis shows survival of silkworm after infection experiment, and the horizontal axis shows dose of (pre -)injected samples (i.e. gelatin or BSA). Each plot (closed circles: gelatin-injected group; open circles: BSA-injected group) represents result of a groups silkworm (n=5). The ED_50_ of gelatin was 1.7 μg/larva, while that of BSA was greater than 250 μg/larva.

I also monitored the acquisition of infection resistance over time (Fig. 2). The complete infection resistance (i.e. 100% survival) to O-157 was observed within 120 minutes after injecting 5.0μg/larva of gelatin (Fig. 2). Given that induction of antimicrobial peptides into hemocoel takes 12 hours after immune priming (Miyashita et al., 2014), it seems unlikely that antimicrobial peptides play major role in the fast immune priming by gelatin components (i.e. collagens).

**Fig. 2.**
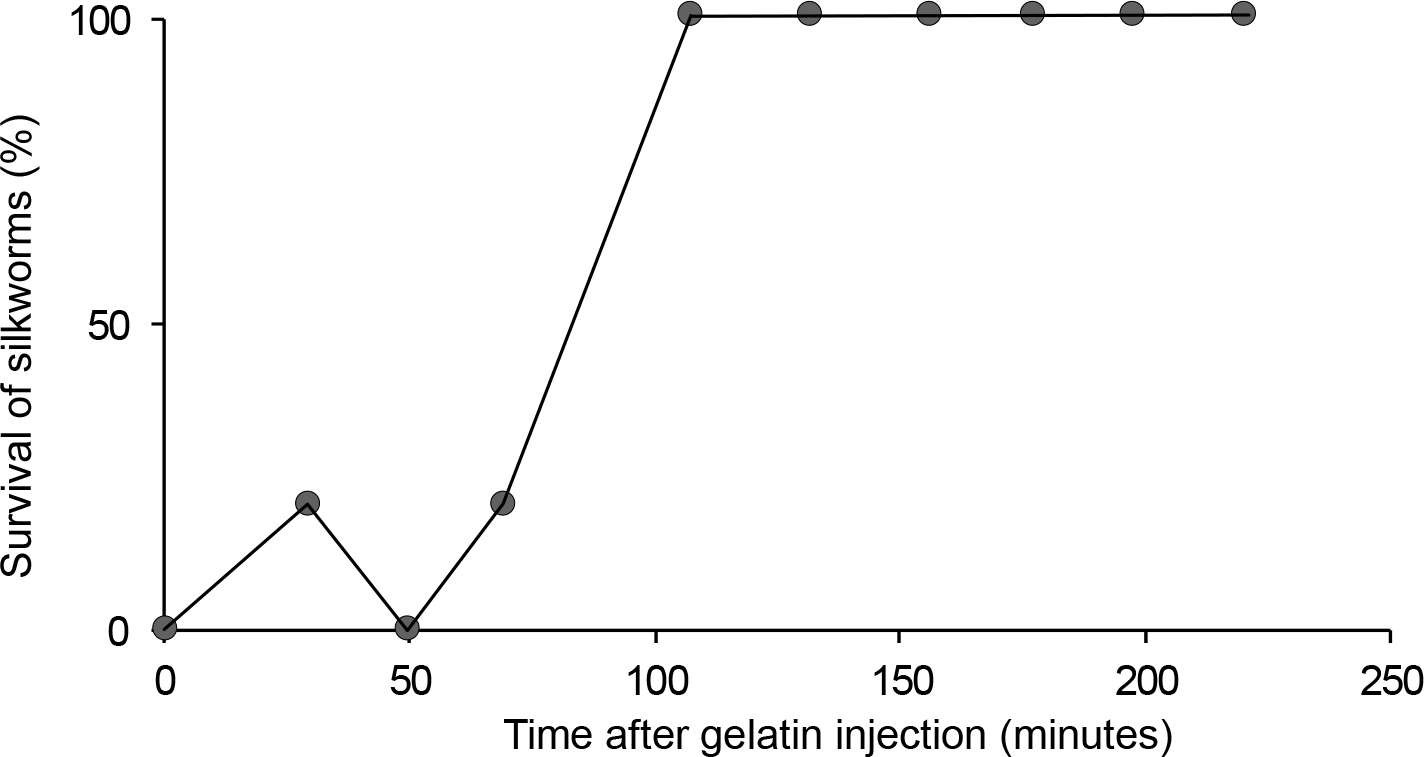
Time course of immune priming triggered by gelatin. The horizontal axis shows time post gelatin injection, where live cells of a pathogenic *E. coli* (O-157:H7) were injected as previously described (Miyashita et al., 2012). The vertical axis indicates survival of silkworm after infection experiment. For each plot, a group of silkworms (n=5) was injected 5 μg/larva of gelatin,incubated at 27Â°C for the given time,and injected the pathogenic *E.coli* cells. Survivals were observed at 20 hours after E. coli injection. The fifth time point is at 110 minutes after an gelatin injection, showing 100% viability in the infection experiment.

Collagens are abundant molecules in connective tissues. Immunological reactions to collagen molecules may have a merit, because, by means of such reaction, animals can prepare in advance for potential infection risks when they get wounds on their exoskeletons or internal tissues (e.g. by predator bites). This is in accordance with ‘Danger’ model (Matzinger, 1994), assuming that collagens are immunoreactive. A recent study demonstrated that a type of collagen is also expressed in hemocytes of silkworm (*B. mori*) and that the hemocytes and fat body cells secretes collagens into hemocoel (Adachi et al., 2005). In the same study, it was also suggested that collagens are present in sheaths formed in encapsulation reaction by hemocytes (Adachi et al., 2005). Given these facts and my finings in this paper, it seems likely that endogenous collagens (either those spread passively from damages tissues, and those secreted actively from immune cells (i.e. hemocytes and fat body cells)) leads to augmentation of immune activation at systemic level. The gelatin/collagen inducible pathway may be one of the missing pieces in insect immunity, and studies to identify receptors or signaling pathways are worth investigating.

## 4. Conclusion

The present study suggests 1) that the silkworm possesses an immune priming pathway that can be triggered by collagen, but not by non-collagen proteins, and 2) that the collagen-inducible pathway is independent from or bypasses the peptidoglycan-inducible immune priming pathway.

## Acknowledgement

This work was supported by JSPS grant #13J08664. I thank Shelley Adamo, Hayato Kizaki, Chikara Kaito, Tatsuhiko Kyuma, and Masaki Ishii for insightful comments and suggestions.

---

Figure S1. Immune priming effect of porcine or bovine collagens in silkworm Silkworms were first injected saline (open circles), 50μg/larva of porcine collagen type I (closed squares), or 50μg/larva of bovine collagen type I (closed triangles). The silkworms were then incubated at 27°C for six hours, and injected live cells of Pseudomonas aeruginosa PAO1 strain. The bacteria was previously shown to be pathogenic on silkworms (Miyashita et al., 2015). The horizontal axis indicates infection dose of *P. aeruginosa* (log-scaled), and the vertical axis indicates survival of silkworm (%) at 12 hours after infection. Each plot represents data from a group of silkworms (n=4).

